# Transcranial alternating current stimulation affects several alpha components depending on their frequencies relative to the stimulation frequency

**DOI:** 10.1101/2023.02.08.527578

**Authors:** Shuka Shibusawa, Tomoya Kawashima, Kaoru Amano

## Abstract

**Background:** The aftereffects of transcranial alternating current stimulation (tACS), especially targeting occipital alpha oscillations, have been reported to show large individual differences in behavioural effects and neural responsiveness. We predicted that this variance at least partly originates from the fact that multiple alpha components are affected differently by alpha-tACS.

**Objective:** To test the above prediction, we decomposed several alpha components from the data and evaluated the aftereffects separately for each component and participant. More specifically, we tested how the difference between the stimulation frequency and peak frequency influences the aftereffects of tACS on each alpha component.

**Methods:** Eighteen participants received 20-min tACS or sham stimulation on separate days. Ten minutes of magnetoencephalography data were collected before and after stimulation, and spectral analysis was performed with a high-frequency resolution (0.1 Hz) to disentangle different alpha components based on the difference in peak frequencies and spatial patterns. Results: The results revealed three alpha components with slightly different frequencies. tACS increased or decreased the power of these alpha components depending on the relative frequency difference from electrical stimulation. Furthermore, observable components differed among participants, possibly because of anatomical differences in each alpha source.

**Conclusion:** When each alpha component is not analysed separately, the change in overall alpha power will be a combination of decreased or increased components, and inter-individual differences will become larger. Our study highlights the importance of noting the presence of multiple alpha components with relatively small differences in the peak frequency for experimental design and analysis.

## 1. Introduction

Neural oscillations, observed as extracellular potentials with electro-or magnetoencephalography (EEG/MEG), reflect rhythmic synchronisation of cell populations, creating functional networks linked to various brain functions (Buzsaki, 2006; Herrmann et al., 2016). The functional roles of neural oscillations have long been explored by measuring responses to sensory stimuli (Fiebelkorn et al., 2013; Romei et al., 2012) or resting-state activities (Bosboom et al., 2006; Dubovik et al., 2012). Recently, many studies using noninvasive brain stimulation (NIBS) techniques, such as rhythmic transcranial magnetic stimulation (rTMS) and transcranial alternating current stimulation (tACS) tuned to physiological frequencies (Herrmann et al., 2016), have demonstrated distinct effects on perception (Feurra et al., 2011b; Helfrich et al., 2014), cognition (Kasten and Herrmann, 2017; Vosskuhl et al., 2015), memory (Reinhart and Nguyen, 2019), and motor-related processes (Feurra et al., 2011a).

In tACS, the polarity of the electrodes reverses periodically at a specific frequency (Andrea Antal et al., 2008), which is thought to cause alternating hyperpolarisation and depolarisation of the membrane potential, resulting in the firing timing of neurones being entrained into the stimulation frequency (Krause et al., 2019). Many studies have reported increased power in the post-stimulation period compared to the pre-stimulation period (an aftereffect) (Kasten et al., 2016; Kasten et al., 2019; Heiko I. Stecher and Herrmann, 2018; Vossen et al., 2015; Zaehle et al., 2010). However, it has been argued that tACS effects are highly variable and cannot be replicated even within participants (Fekete et al., 2018; Veniero et al., 2017), although the physical amount of stimulation and individual differences in occipital alpha distribution explain some of the individual differences in aftereffects (Kasten et al., 2019). To understand the factors underlying these variabilities and to control the brain oscillations to the desired state for improvement of brain functions, we tested the tACS aftereffects of alpha oscillations (7–13 Hz), as they are the most prominent oscillations in the human brain (Barzegaran et al., 2017).

While alpha oscillations are known to be dominant in the occipital region, they are also observable in other regions, such as the tau rhythm in the temporal region and mu rhythm in the parietal region (Narici et al., 1998; Takahashi and Kitazawa, 2017). It is unknown how tACS, which has a low focality (A. Antal and Paulus, 2013), affects the alpha components in various regions over stimulation areas such as the occipital region. Therefore, we explored the mechanism underlying the aftereffects of tACS by decomposing multiple alpha components with different frequencies and spatial characteristics and evaluating the aftereffects of each component.

## 2. Material and methods

Experiments were conducted in a within-participants design to compare the effects of tACS between the tACS and sham-stimulation conditions (Figure 1A). Before the tACS sessions, to estimate an individual alpha frequency, the spontaneous alpha activities in the occipital area were recorded with eyes open and closed twice for each participant, for a total of four resting-state measurements (2 conditions × 2 repetitions). In the eyes-open condition, participants were instructed to gaze at a fixation point on a display screen in front of them. The peak alpha frequency in the eyes-open condition was used as the individual stimulation frequency (ISF). All recordings were 120 seconds long for each measurement.

**Figure 1.**
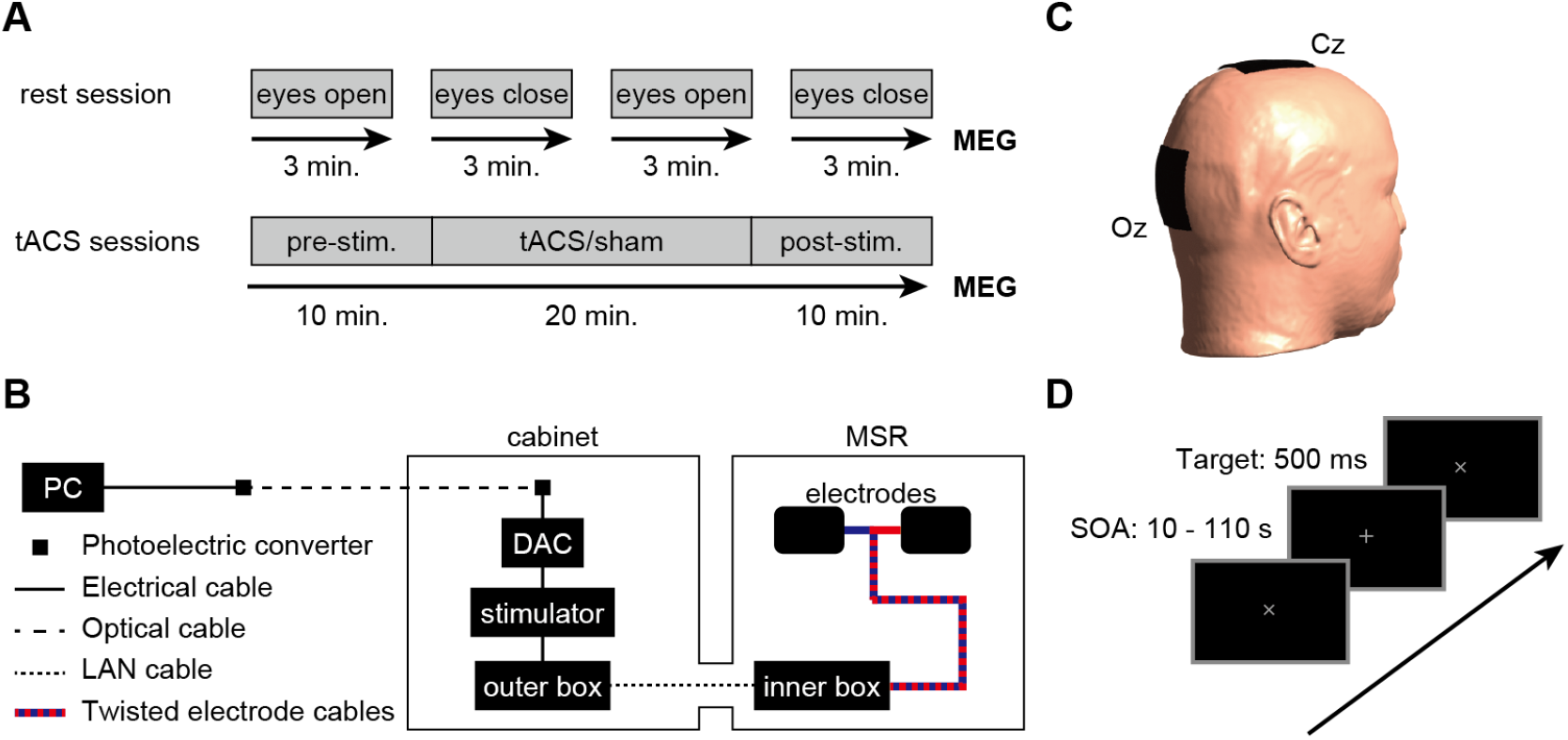
Experimental procedure. (A) Time course of the experiment. On the first day, we conducted the resting-state measurement to estimate the individual alpha frequency to set the individual stimulus frequency (ISF). On the second and third days, all participants received either tACS or sham stimulation. The order of stimulation conditions was counterbalanced across participants. Both experimental sessions were conducted at least four days apart. The tACS or sham stimulation was applied during the middle 20 min of the total 40-minute MEG measurement. No stimulation was applied for 10 min each before and after the stimulation period, which was used to evaluate the alpha power changes. (B) tACS setups. The stimulation waveform was streamed to a battery-driven magnetic resonance-compatible tACS device through a digital/analogue converter (DAC). Both the DAC and stimulator were placed in the stimulation cabinet to prevent external electrical noise due to tACS entering the magnetically shielded room. When sending the stimulation current into the cabinet, it must be converted once outside the cabinet to an optical signal and then reconverted to an electrical signal inside the cabinet. Then, we sent the stimulation current to silicon rubber electrodes via a tube in the wall through a shielded LAN cable and a fully twisted pair of cables. (C) Electrode montage. Stimulation electrodes (5 cm × 7 cm) were attached to the Cz and Oz montage, which can produce the highest current densities in posterior brain regions. (D) Visual vigilance task. Participants were asked to fixate on a grey cross in the centre of a black screen. The fixation cross was rotated by 45° for 500 ms after from 10 to 110 s stimulus onset asynchrony (SOA), and the participants had required to respond by pressing a button with their right index finger.

In the tACS sessions, all participants received tACS or sham stimulation in one out of two experimental sessions. The order of the stimulation conditions was counterbalanced across participants. Both experimental sessions were conducted at least four days apart. MEG measurements were performed for 40 min. tACS or sham stimulation was applied during the middle 20 min of the measurement. No stimulation was applied for 10 min before and after the stimulation period, and these 10 min periods were used to evaluate the changes in alpha power. Participants performed a visual vigilance task to maintain their attention for 40 min.

### 2.1. Participants

Twenty-five healthy participants were recruited for the study. The data from seven were excluded from the analysis because their task performance was too low in one of the sessions (lower than 60% for an easy task, see below), mainly due to the participant falling asleep during the experiment. Here, we report the data from 18 participants (8 females, mean age: 22.4 years, SD: 1.4, age range: 20–26 years, Japanese, all right-handed). They provided written informed consent and received monetary compensation for their participation. The experimental protocol was approved by the ethics committee of the National Institute of Information and Communications Technology.

### 2.2. Magnetoencephalogram

MEG measurements were performed using a 360-channel whole-head MEG system (Elekta Neuromag® 360TM, MEGIN/Elekta Oy, Helsinki, Finland) comprising 204 planar gradiometers, 102 magnetometers, and 54 tangential sensors (36 gradiometers and 18 magnetometers) placed in a magnetically shielded room (MSR). In the resting session, the magnetic signals were recorded at 5 kHz in the upright sitting position (60° dewar orientation) in darkness, and a hardware low-pass filter of 1,660 Hz and a high-pass filter of 0.03 Hz were applied online. During the tACS sessions, measurements were performed in the supine position to minimise head movement. The magnetic signals were recorded at 1 kHz in a dimly lit MSR to avoid an increase in the alpha power during the stimulation period (H. I. Stecher et al., 2017). A hardware low-pass filter (330 Hz) and high-pass filter (0.03 Hz) were applied online.

### 2.3. tACS setup

The digital signal of the stimulation waveform, at a sampling rate of 10 kHz, was streamed to a digital/analogue converter (DAC; USB-6211, National Instruments, Austin, TX, USA) using the data acquisition toolbox provided in MATLAB 2017a (MathWorks Inc., Natick, MA, USA). The analogue output from the DAC was connected to the remote input of a battery-driven magnetic resonance-compatible tACS device (DC Stimulator Plus; neuroConn GmbH, Ilmenau, Germany). To prevent external electrical noise from the tACS entering the MSR, both the DAC and the stimulator were placed in the stimulation cabinet. To send the stimulation signal inside the cabinet, it must be converted to an optical signal outside the cabinet, and then reconverted to an electrical signal inside the cabinet (Figure 1B). The stimulator output signal was sent to a connector box located inside the MSR via a tube in the wall through a shielded cable, from which a fully twisted pair of cables delivered current to the silicon rubber stimulation electrodes (Herring et al., 2019). Stimulation electrodes (5 cm × 7 cm) were attached to the Cz and Oz according to the 10-20 system (Figure 1C) using a conductive paste (Ten20, Weaver & Co., Aurora, CO, USA). This electrode montage was used because a previous modelling study showed that it produced the highest current densities in posterior brain regions (Neuling et al., 2012). The electrode impedance was maintained below 20 kΩ (including 10 kΩ resistors in the magnetic resonance-compatible stimulator cables).

Following a 10-min no-stimulation period, tACS or sham stimulation was applied for 20 min at the ISF determined from resting session recordings. In the tACS condition, stimulation was applied with an intensity of 1.3 mA (peak-to-peak) and 4 s fade-in and fade-out intervals at the beginning and end of the stimulation period, respectively. In the sham-stimulation condition, 8 s of stimulation (4 s of fade-in and 4 s of fade-out) was applied before and after the stimulation interval, with no stimulation in the stimulation interval.

### 2.4. Visual vigilance task

To ensure that the participants maintained their arousal and attention to the task for 40 min, they were asked to perform a visual vigilance task as used by Kasten et al. (2019). Visual stimuli were generated using Psychtoolbox 3 (Kleiner et al., 2007) working on MATLAB and were presented on a screen in an MSR using an LCD projector (PT-DZ680, Panasonic, Osaka, Japan; 60 Hz refresh rate) approximately 60 cm from the participant.

A grey fixation cross with a 1.5° visual angle was presented on a black background at the canter of the screen. Participants were asked to press a button with their right index finger when they detected a rotation which randomly occurred for 500 ms (Figure 1D). The stimulus onset asynchrony (SOA) was set from 10 to 110 s. Only trials in which the button was pressed within 2 s after starting the rotation were defined as successful trials, and only participants with a daily task performance higher than 60% were used for later analysis (median accuracy = 90%, *min* = 24%, *max* = 100%).

### 2.5. MEG preprocessing

We first applied the oversampled temporal projection (OTP), an uncorrelated sensor noise suppression algorithm, which works with any multichannel measurement system that has sufficient sensor density to spatially oversample the signal of interest (Larson and Taulu, 2018). Second, bad channels were automatically identified using the Xscan software version 3.0.17 (MEGIN/Elekta Oy, Helsinki, Finland). Third, the detected bad channels were substituted with interpolated values, and external interference in the MEG was suppressed by signal space separation (SSS) (Taulu et al., 2005) using Maxfilter-2.2TM (MEGIN/Elekta Oy, Helsinki, Finland) for the resting-session data. For the tACS session data, after removing 20 min of data during tACS, external interferences of 10 min of pre- and post-stimulation data were suppressed using temporal signal space separation (tSSS) instead of SSS. This is because tACS electrodes might cause temporally correlated noise over a large area. For the tSSS parameters, the correlation limit was set to 0.998, which was higher than the default value because of the use of OTP (Clarke et al., 2020). All other settings were the default.

### 2.6. Determination of tACS frequency

All data were analysed using the MNE-Python toolbox version 19.0 (Gramfort et al., 2013). The signals during the resting session were band-pass filtered between 1 Hz and 50 Hz. Then, artefacts reflecting heartbeats or blinks were removed using the FastICA algorithm. Continuous signals were then cut into 2 s segments. Segments containing evident noise were excluded by visual inspection. Power spectral densities (PSDs) were then calculated using a multitaper fast Fourier transform (FFT) on all segments (8 s zero padding, apparent frequency resolution = 0.1 Hz). Finally, the PSDs of the eight occipital gradiometers (MEG2012, MEG2013, MEG2022, MEG2023, MEG2032, MEG2033, MEG2042, and MEG2043 in the Neuromag MEG system) in the eyes-open condition data were averaged, and its peak frequency within the alpha band (7–13 Hz) was set to be the ISF.

### 2.7. Analysis of the tACS effects

The pre-processed data were down-sampled to 250 Hz. The signals were then band-pass filtered between 1 and 40 Hz. Subsequently, the components reflecting heartbeats or blinks were removed using the FastICA algorithm. After removing the artefacts, continuous signals were cut into 10 s segments. Segments containing the timing of the rotation of the visual stimuli and button responses were excluded. Segments containing apparent noise were excluded by visual inspection. PSDs were calculated using a multitaper FFT (frequency resolution = 0.1 Hz) for the first 200 artefact-free segments of each period.

To focus on neural oscillations without non-oscillatory activities, exponential approximation and Gaussian fitting were used to parameterise periodic and aperiodic components using the fitting-oscillations-and-one-over-f (FOOOF) toolbox version 1.0.0 (https://fooof-tools.github.io/) (Donoghue et al., 2020). FOOOF was applied to the averaged PSDs across all gradiometers (detailed parameter settings are provided in Supplementary Information 1), and the aperiodic components were subtracted for the following analysis. Two prominent peaks were visible in most participants in the alpha frequency range (individual fitting results are shown in Supplementary Figure 1). To consider not only the peaks visible from the original spectrum but also those that are not necessarily visible but are detected by Gaussian fitting, we multiplied the height of the spectrum of the original signals by the height of the fitted Gaussian distribution. The two main components for each participant were defined as the two frequencies with the largest multiplied values from 7.0 to 13.0 Hz. The 204 gradiometers used in this analysis were located at 102 locations, two at each location in different orientations; thus, the root mean square of the PSDs of the two sensors at each location was calculated. Then, to compensate for the distance-dependent absolute power values, the sum of the power around the observed peaks (± 0.3 Hz) was normalised by dividing by the total power at 1–40 Hz. Topographic maps were plotted with relative power values, with the minimum and maximum values set to 0 and 1, respectively (Figure 2).

**Figure 2.**
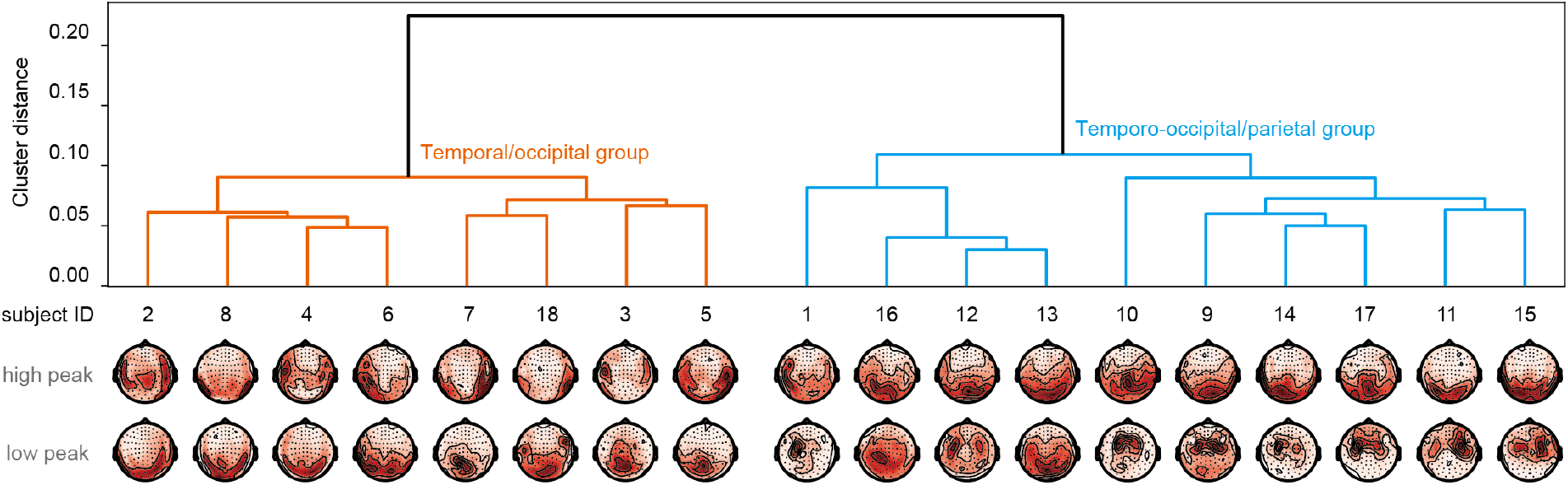
Two groups of participants obtained by cluster analysis based on their topographic maps. Cluster analysis revealed that the spatial distributions of the two frequency peaks obtained by frequency analysis with a high-frequency resolution were divided into two groups, one with temporal and occipital peaks and the other with temporo-occipital and parietal peaks. We evaluated the changes in the power of multiple alpha components from pre-to post-stimulation in these two groups separately.

## 3. Results

### 3.1. Performance of the visual vigilance task

The average performance of the visual vigilance task was *M* = 88.5% ± *SD* = 12.5% in the sham-stimulation condition and *M* = 91.8% ± *SD* = 7.7% in the tACS condition. The stimulation conditions did not cause a significant difference in task performance (*t* (34) = 1.05, *p* = 0.31, Cohen’s *d* = 0.24, paired *t*-test).

### 3.2. Separation into two groups of participants based on FFT with a high-frequency resolution

Frequency analysis of the MEG data pre- and post-stimulation with a high-frequency resolution (0.1 Hz) revealed multiple peaks in the alpha band (spectrum post-stimulation for sham and stimulation conditions are shown in Supplementary Figure 1A and B, respectively; the results obtained by FFT with a low-frequency resolution are shown in Supplementary Information 2). These peaks corresponded to the occipital and temporal components for some participants, while they corresponded to the occipital and parietal components for other participants (Barzegaran et al., 2017). To group the participants based on peak spatial patterns, we performed cluster analysis on the two largest peaks (see Supplementary Information 3). Individual topographic maps showed that in eight of the 18 participants, lower and higher peaks appeared in the temporal and occipital regions, respectively. In the remaining ten participants, lower and higher peaks appeared in the temporo-occipital and parietal regions, respectively (Figure 2). These two peaks were identified regardless of the stimulation condition for each participant (Supplementary Figure 1). Therefore, based on these stable characteristics of the alpha components, we termed them the temporal/occipital and temporo-occipital/parietal groups.

### 3.3. tACS at the alpha frequency affects multiple alpha components differently

#### 3.3.1. Temporal/occipital group

Here, we examined the changes in alpha power from pre-to post-stimulation in each group of participants. We first analysed the power of alpha peaks of the temporal/occipital group using a three-way repeated-measures analysis of variance (ANOVA) with factors of 2 (alpha component: high-frequency component, low-frequency component) × 2 (stimulation: tACS, sham stimulation) × 2 (time: pre- and post-stimulation). As a result, the three-way interaction of the alpha component, simulation, and time was significant (*F* (1, 7) = 25.69, *p* = .001,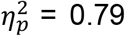); therefore, we performed 2 (stimulation) × 2 (time) repeated-measures ANOVAs for each alpha component.

In the temporal component of the temporal/occipital group (Figure 3A, left), the main effect was significant for the time factor (*F* (1, 7) = 6.42, *p* = .039, 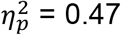), but not for the stimulation factor (*F* (1, 7) = 1.00, *p* = .351,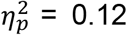). Their interactions tended to be significant (*F* (1, 7) = 4.63, *p* = .068, 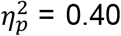). These analyses suggest that the temporal component of the temporal/occipital group showed a power increase, regardless of the stimulus condition. Thus, the increase tended to be slightly weaker in the tACS condition than in the sham-stimulation condition.

**Figure 3.**
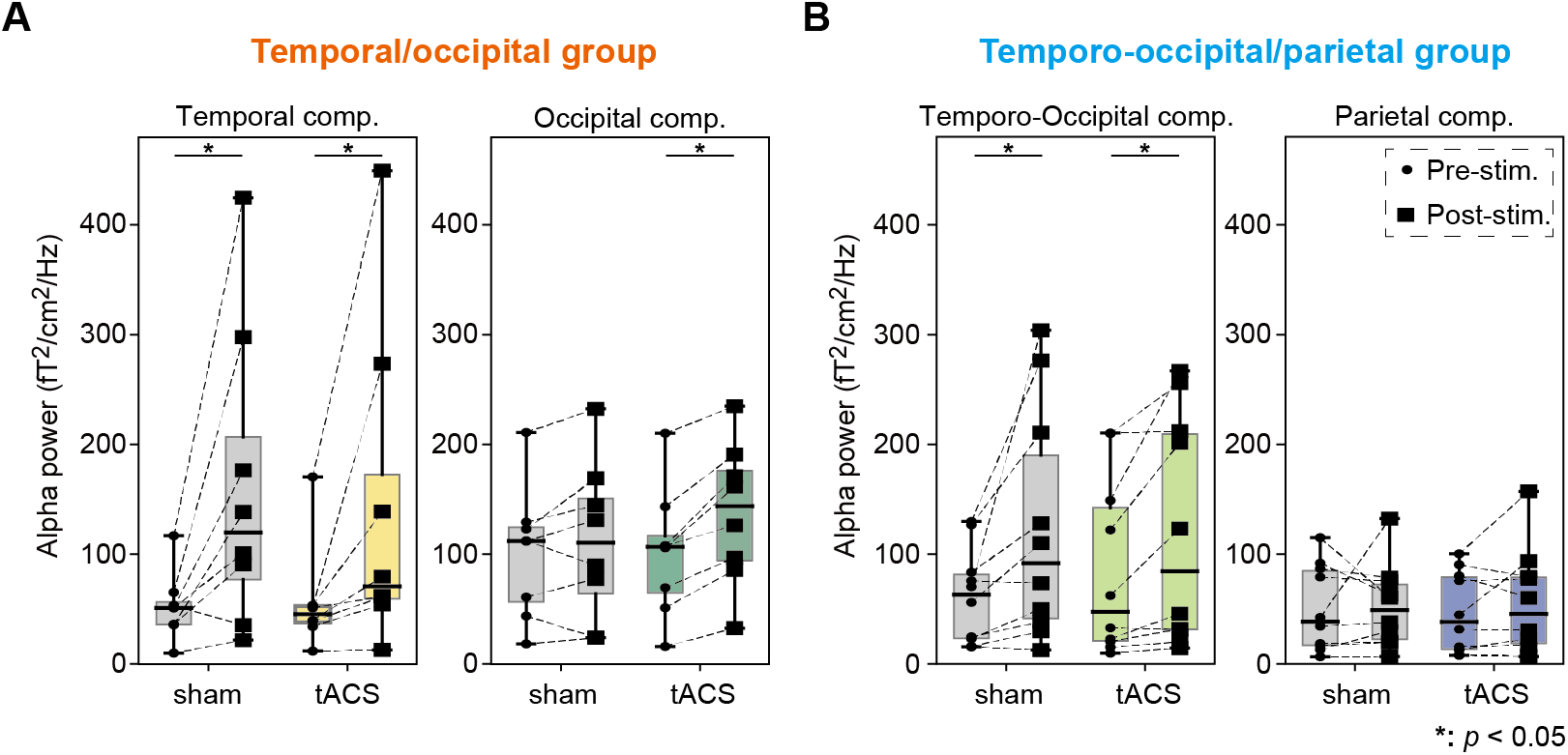
The power changes from pre-to post-stimulation of each alpha component. (A) The results of the temporal/occipital group. The temporal alpha power was increased in both stimulation conditions, and its increase tended to be suppressed in the tACS condition. The occipital alpha power was increased only in the tACS condition. (B) The results of the temporo-occipital/parietal group. The temporo-occipital alpha power was increased in both stimulation conditions. No significant changes were observed in the parietal component.

In the occipital component of the temporal/occipital group (Figure 3A right), a two-way repeated-measures ANOVA revealed a significant main effect of the time factor (*F* (1, 7) = 17.92, *p* = .004, 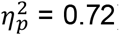), but not of the stimulation factor (*F* (1, 7) = 2.12, *p* = .189, 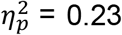). A significant interaction was also observed (*F* (1, 7) = 7.86, *p* = .026, 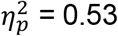). Further analysis of the simple main effects revealed that the power of the occipital component in the post-stimulation period was significantly higher than that in the pre-stimulation period in the tACS condition (*F* (1, 7) = 33.72, *p* = .001, 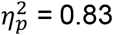), but not in the sham-stimulation condition (*F* (1, 7) = 1.67, *p* = .237, 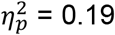). In other words, the power of the occipital component in the temporal/occipital group was increased by the tACS.

#### 3.3.2. Temporo-occipital/parietal group

A three-way repeated-measures ANOVA on the temporo-occipital/parietal group revealed a significant main effect in the alpha component factor (*F* (1, 9) = 5.55, *p* = .043, 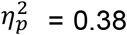), but not in the time and stimulation factors (*F* (1, 9) = 4.78, *p* = .057, 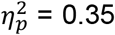; *F* (1, 9) = 0.22, *p* = .650, 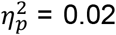). The two-way interaction between the alpha component and time was significant (*F* (1, 9) = 11.56, *p* = .008, 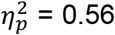). In contrast, the interactions of the alpha component and stimulation and stimulation and time were not significant (*F* (1, 9) = 0.80, *p* = .395, 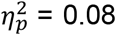; *F* (1, 9) = 0.83, *p* = .385, 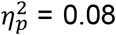). The three-way interaction was not significant (*F* (1, 9) = 4.23, *p* = .070, 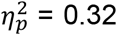). Further analysis of the simple main effects revealed that the power of the temporo-occipital component in the post-stimulation period was significantly higher than that in the pre-stimulation period, regardless of the condition (*F* (1, 9) = 7.76, *p* = .021, 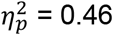; Figure 3B, left), whereas the power of the parietal component was unchanged (*F* (1, 9) = 0.35, *p* = .568, 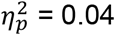; Figure 3B, right).

## 4. Discussion

In this study, we attempted to provide insight into the factors underlying the interparticipant variability of tACS aftereffects. To this end, we decomposed multiple alpha components and evaluated the power change from pre-to post-tACS for each alpha component separately. Decomposition of the alpha components revealed that there were two groups of participants that showed different alpha patterns: a group with dominant temporal and occipital components, and a group with dominant temporo-occipital and parietal components. By normalising the power change of each component with the changes between pre- and post-stimulation in the sham-stimulation condition, the tACS effects can be expressed as follows (Figure 4). In the temporal/occipital group, the temporal alpha power tended to decrease, whereas the occipital alpha power increased due to tACS. The temporo-occipital/parietal group, temporal-occipital and parietal alpha power did not show any significant differences between the stimulation conditions. Here, we discuss the effect of tACS on each component considering the frequency mismatch between each alpha component and tACS.

**Figure 4.**
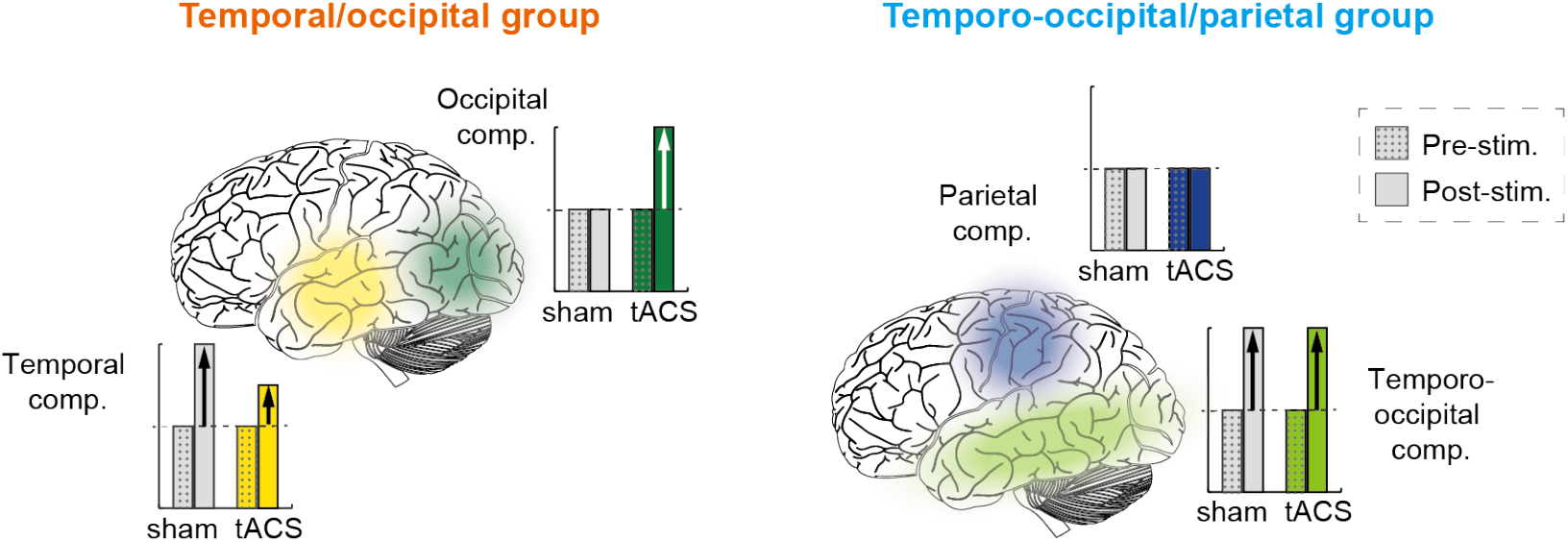
A schematic illustration of the tACS aftereffects on each alpha component. The tACS decreased the temporal alpha power and increased the occipital alpha power in the temporal/occipital group, while did not affect either component of the temporo-occipital/parietal group.

### 4.1. Dominant alpha patterns differ among participants

In frequency analyses, the time and frequency resolutions have a trade-off relationship. As most of the previous studies addressing alpha oscillations were analysed using a short time length of approximately 1–2 s (Kasten et al., 2016; Kasten et al., 2019; Heiko I. Stecher and Herrmann, 2018; Vossen et al., 2015), a single peak appears in the alpha band, which is conventionally defined as an individual alpha frequency. However, the temporal tau and parietal mu rhythms are also known to be approximately10 Hz, whose frequencies are slightly lower and higher than the occipital alpha rhythms, respectively (Yokosawa et al., 2020). It can therefore be expected that multiple components are contained in an apparent single peak because the measured brain signals include spatially mixed components due to volume conduction (Benwell et al., 2019).

As this study did not focus on the fast temporal dynamics of alpha oscillations, it was possible to increase the length of the time window for frequency analysis, thereby increasing the frequency resolution. We successfully identified multiple peaks for each participant (Supplementary Figure 1). Three peaks corresponding to occipital, parietal, and temporal components were observed, but not necessarily all from each participant. First, no parietal peaks were obtained for some participants (i.e., the temporal/occipital group). This is consistent with previous reports that parietal mu rhythms were only observed in approximately one-half to one-third of participants (Kuhlman, 1978; Vanni et al., 1999). Second, the temporal and occipital components could not be separated in other participants (i.e., the temporo-occipital/parietal group). The temporal tau component has been suggested to be observed only in some participants, possibly because the orientation of the dipole differs depending on the participant’s brain structure, making the sensor less sensitive. It is also possible that increased occipital alpha power, depending on the participant’s state, masks the temporal tau rhythm (Yokosawa et al., 2020). In the present results, a single peak extended over both the temporal and occipital regions, so it is reasonable to understand that the temporal and occipital components overlap in the temporo-occipital/parietal group.

As described above, the observation that the detectable alpha pattern differs among participants even when the frequency resolution is increased can be attributed to individual differences rather than a limitation of the analysis method. Therefore, in this study, it would be reasonable to group participants using clustering analysis to evaluate the effects of tACS on each component.

### 4.2. Alpha power changes of each component due to tACS

In the temporal/occipital group, occipital alpha was significantly increased by tACS, while temporal alpha was reduced (Figure 4). These effects could be explained by the difference between the frequency of each alpha component and stimulation frequency. As shown in Figure 5A, the frequency of the occipital component was slightly slower than that of the ISF, whereas that of the temporal component was slightly faster than that of the ISF.

**Figure 5.**
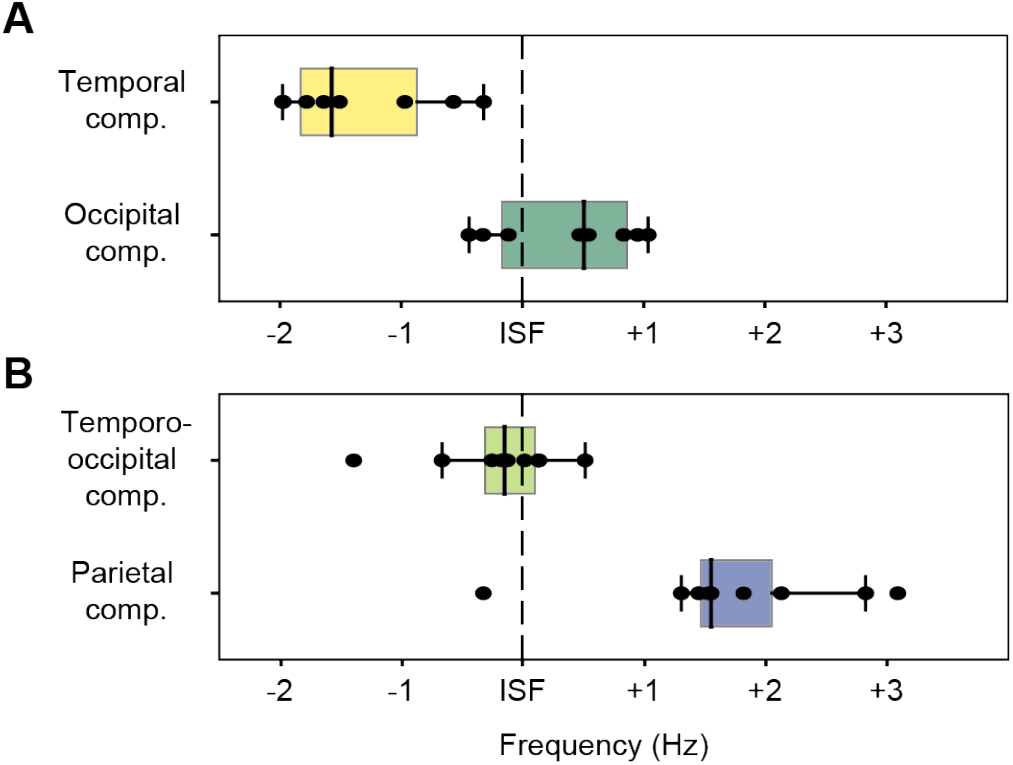
Frequency mismatches between each alpha component and the individual stimulation frequency (ISF). (A) In the temporal/occipital group, the frequency of the temporal component was slower, and that of the occipital component was faster than the ISF. These mismatches may cause long-term potentiation (LTP) or depression (LTD). (B) In the temporo-Occipital/Parietal group, because the frequency of the temporo-occipital component of each participant was faster or slower than the stimulation frequency, it is possible that LTP and LTD occurred depending on the participants, and the effects were cancelled out. A much faster frequency of the parietal component would have led to a lack of the tACS effect.

Based on these frequency mismatches, we suggest that spike-timing-dependent plasticity (STDP) (Fröhlich, 2016) could explain our results as follows. Firs, the mechanism of neural oscillation formation consists of two parts (Vossen et al., 2015): the generator of the electrical potential and the recurrent loop that generates oscillatory potential with the frequency of each alpha component (e.g., occipital, temporal, and parietal alpha). tACS stimulates this generator at a specific frequency and entrains it into the stimulation frequency (Johnson et al., 2020; Krause et al., 2019). When the entrained frequency (i.e., stimulation frequency) is slightly slower than the frequency of that loop (i.e., the frequency of each alpha component), long-term potentiation (LTP) occurs due to timing-dependent firing over several tens of minutes, resulting in an increase in the power of the loop frequency (Elyamany et al., 2021; Vossen et al., 2015; Zaehle et al., 2010). This is exactly what we found for occipital alpha. On the other hand, when the entrained frequency is slightly faster than the frequency of that loop, long-term depression (LTD) occurs, resulting in a decrease in the power of the loop frequency. This is what we found for temporal alpha.

The temporal alpha power of the temporal/occipital group tended to decrease with applying tACS. One possible reason for the effect of occipital tACS on temporal alpha is that the tACS currents spread not only to the area directly under the electrodes but also to surrounding areas, such as the temporal region, resulting in it being affected by direct stimulation. The other possibility is that the loops of each alpha component are connected and indirectly affected by the directly entrained component, as is observed for occipital alpha entrainment by auditory stimuli (Bauer et al., 2020; Bauer et al., 2021). Since the number of participants in the temporal/occipital group was small and the statistical power was weak, the effect of tACS on nontarget areas requires further investigation.

In the temporo-occipital/parietal group, neither the temporo-occipital nor parietal component showed significant changes due to tACS (Figure 4). These results can also be explained by STDP in the same manner as described above. At the temporo-occipital component, whose frequency of each participant was faster or slower than the stimulation frequency (Figure 5B), LTP or LTD occurred depending on the participant, and the effects were cancelled out. The lack of the tACS effect for the parietal component was probably due to its much faster frequency than the stimulation frequency (Figure 5B). A study using rat hippocampal neurones indicated that LTP or LTD occurs when repetitive postsynaptic spiking occurs within a relatively short interval after or before presynaptic activation (Bi and Poo, 1998).

### 4.3. Temporal alpha power increase due to drowsiness

Temporal alpha power significantly increased even in the sham-stimulation condition (Figure 3). As shown in Supplementary Table 1, these components are relatively slow in the alpha band and are generally referred to as low alpha (Kubicki et al., 1979). The power of this low alpha is known to increase with time (Benwell et al., 2019), possibly due to mental fatigue or drowsiness (Boksem et al., 2005; Craig et al., 2012; Simon et al., 2011). In the present study, the participants were asked to perform a visual vigilance task for 40 min, which possibly induced fatigue or drowsiness, and increased the temporal alpha power. In fact, in the subjective evaluation of drowsiness, more than half of the participants chose “I became sleepy towards the second half of the session” or “I was sleepy the whole time” (See Supplementary Information 4 and Supplementary Figure 3 for further details).

## 5. Conclusion

Our results demonstrate that tACS show different effects on multiple alpha components depending on the mismatch between the frequency of the component and the stimulation frequency. Namely, even if the stimulation is performed to enhance power, it may be conversely suppressed depending on the stimulation frequency. These opposite effects will increase inter-individual differences and may be the reason why previous studies demonstrated inconsistent results (Fekete et al., 2018; Veniero et al., 2017). In the tACS experiment, the stimulation frequency should be carefully set by considering these characteristics. In addition, in the analysis, each component should be evaluated separately by increasing the frequency resolution. This is because if LTP in one component and LTD in another are mixed in the evaluation, the power change will be misinterpreted. Furthermore, if multiple peaks are analysed as a single peak, the change in the power balance between the multiple peaks can be incorrectly associated with the change in the frequency of a single peak (Supplementary Figure 2). To summarise, for the tACS experiments on alpha oscillations, we should note the existence of several alpha components with relatively small differences in peak frequency, not only for the experimental design but also for the analysis.

## Acknowledgement

Funding: This work was supported by the Japan Society for the Promotion of Science (JSPS) Grant-in-Aid for JSPS Fellows (grant number 19J22132) to S.S.; JSPS Grant-in-Aid for Early-Career Scientists (grant number JP20K14274) to T.K.; JSPS Grant-in-Aid for Scientific Research on Innovative Areas (grant number 18H05523) to K.A.; and JSPS Grant-in-Aid for Scientific Research (A) (grant number 21H04909) to K.A. We would like to thank Editage (www.editage.com) for English editing.

## References

Antal, A., Boros, K., Poreisz, C., Chaieb, L., Terney, D., Paulus, W., 2008. Comparatively weak after-effects of transcranial alternating current stimulation (tACS) on cortical excitability in humans. Brain Stimul. 1(2), 97–105. doi:https://doi.org/10.1016/j.brs.2007.10.001

Antal, A., Paulus, W., 2013. Transcranial alternating current stimulation (tACS). Front. Hum. Neurosci. 7, 317. doi:https://doi.org/10.3389/fnhum.2013.00317

Barzegaran, E., Vildavski, V. Y., Knyazeva, M. G., 2017. Fine Structure of Posterior Alpha Rhythm in Human EEG: Frequency Components, Their Cortical Sources, and Temporal Behavior. Sci. Rep. 7(1), 8249. doi:https://doi.org/10.1038/s41598-017-08421-z

Bauer, A.-K. R., Debener, S., Nobre, A. C., 2020. Synchronisation of Neural Oscillations and Cross-modal Influences. Trends Cogn. Sci. 24(6), 481–495. doi:https://doi.org/10.1016/j.tics.2020.03.003

Bauer, A.-K. R., van Ede, F., Quinn, A. J., Nobre, A. C., 2021. Rhythmic Modulation of Visual Perception by Continuous Rhythmic Auditory Stimulation. The Journal of Neuroscience 41(33), 7065. doi:https://doi.org/10.1523/JNEUROSCI.2980-20.2021

Benwell, C. S. Y., London, R. E., Tagliabue, C. F., Veniero, D., Gross, J., Keitel, C., Thut, G., 2019. Frequency and power of human alpha oscillations drift systematically with time-on-task. NeuroImage 192, 101–114. doi:https://doi.org/10.1016/j.neuroimage.2019.02.067

Bi, G.-q., Poo, M.-m., 1998. Synaptic Modifications in Cultured Hippocampal Neurons: Dependence on Spike Timing, Synaptic Strength, and Postsynaptic Cell Type. The Journal of Neuroscience 18(24), 10464–10472. doi:https://doi.org/10.1523/jneurosci.18-24-10464.1998

Boksem, M. A. S., Meijman, T. F., Lorist, M. M., 2005. Effects of mental fatigue on attention: An ERP study. Cognitive Brain Research 25(1), 107–116. doi:https://doi.org/10.1016/j.cogbrainres.2005.04.011

Bosboom, J. L. W., Stoffers, D., Stam, C. J., van Dijk, B. W., Verbunt, J., Berendse, H. W., Wolters, E. C., 2006. Resting state oscillatory brain dynamics in Parkinson’s disease: An MEG study. Clin. Neurophysiol. 117(11), 2521–2531. doi:https://doi.org/10.1016/j.clinph.2006.06.720

Buzsaki, G., 2006. Rhythms of the Brain: Oxford university press.

Clarke, M., Larson, E., Tavabi, K., Taulu, S., 2020. Effectively combining temporal projection noise suppression methods in magnetoencephalography. J. Neurosci. Methods 341, 108700. doi:https://doi.org/10.1016/j.jneumeth.2020.108700

Craig, A., Tran, Y., Wijesuriya, N., Nguyen, H., 2012. Regional brain wave activity changes associated with fatigue. Psychophysiology 49(4), 574–582. doi:https://doi.org/10.1111/j.1469-8986.2011.01329.x

Donoghue, T., Haller, M., Peterson, E. J., Varma, P., Sebastian, P., Gao, R., … Voytek, B., 2020. Parameterizing neural power spectra into periodic and aperiodic components. Nat. Neurosci. 23(12), 1655–1665. doi:https://doi.org/10.1038/s41593-020-00744-x

Dubovik, S., Pignat, J.-M., Ptak, R., Aboulafia, T., Allet, L., Gillabert, N., … Guggisberg, A. G., 2012. The behavioral significance of coherent resting-state oscillations after stroke. NeuroImage 61(1), 249–257. doi:https://doi.org/10.1016/j.neuroimage.2012.03.024

Elyamany, O., Leicht, G., Herrmann, C. S., Mulert, C., 2021. Transcranial alternating current stimulation (tACS): from basic mechanisms towards first applications in psychiatry. Eur. Arch. Psychiatry Clin. Neurosci. 271(1), 135–156. doi:https://doi.org/10.1007/s00406-020-01209-9

Fekete, T., Nikolaev, A. R., De Knijf, F., Zharikova, A., van Leeuwen, C., 2018. Multi-Electrode Alpha tACS During Varying Background Tasks Fails to Modulate Subsequent Alpha Power. Front. Neurosci. 12, 428. doi:https://doi.org/10.3389/fnins.2018.00428

Feurra, M., Bianco, G., Santarnecchi, E., Del Testa, M., Rossi, A., Rossi, S., 2011a. Frequency-Dependent Tuning of the Human Motor System Induced by Transcranial Oscillatory Potentials. The Journal of Neuroscience 31(34), 12165. doi:https://doi.org/10.1523/JNEUROSCI.0978-11.2011

Feurra, M., Paulus, W., Walsh, V., Kanai, R., 2011b. Frequency specific modulation of human somatosensory cortex. Front. Psychol. 2, 13. doi:https://doi.org/10.3389/fpsyg.2011.00013

Fiebelkorn, I. C., Snyder, A. C., Mercier, M. R., Butler, J. S., Molholm, S., Foxe, J. J., 2013. Cortical cross-frequency coupling predicts perceptual outcomes. NeuroImage 69, 126–137. doi:https://doi.org/10.1016/j.neuroimage.2012.11.021

Fröhlich, F. 2016. Chapter 4 - Synaptic Plasticity. In F. Fröhlich (Ed.), Network Neuroscience (pp. 47–58). San Diego: Academic Press.

Gramfort, A., Luessi, M., Larson, E., Engemann, D., Strohmeier, D., Brodbeck, C., … Hämäläinen, M., 2013. MEG and EEG data analysis with MNE-Python. Front. Neurosci. 7(267). doi:https://doi.org/10.3389/fnins.2013.00267

Helfrich, Randolph F., Schneider, Till R., Rach, S., Trautmann-Lengsfeld, Sina A., Engel, Andreas K., Herrmann, Christoph S., 2014. Entrainment of Brain Oscillations by Transcranial Alternating Current Stimulation. Curr. Biol. 24(3), 333–339. doi:https://doi.org/10.1016/j.cub.2013.12.041

Herring, J. D., Esterer, S., Marshall, T. R., Jensen, O., Bergmann, T. O., 2019. Low-frequency alternating current stimulation rhythmically suppresses gamma-band oscillations and impairs perceptual performance. NeuroImage 184, 440–449. doi:https://doi.org/10.1016/j.neuroimage.2018.09.047

Herrmann, C. S., Strüber, D., Helfrich, R. F., Engel, A. K., 2016. EEG oscillations: From correlation to causality. Int. J. Psychophysiol. 103, 12–21. doi:https://doi.org/10.1016/j.ijpsycho.2015.02.003

Johnson, L., Alekseichuk, I., Krieg, J., Doyle, A., Yu, Y., Vitek, J., … Opitz, A., 2020. Dose-dependent effects of transcranial alternating current stimulation on spike timing in awake nonhuman primates. Science Advances 6(36), eaaz2747. doi:https://doi.org/10.1126/sciadv.aaz2747

Kasten, F. H., Dowsett, J., Herrmann, C. S., 2016. Sustained Aftereffect of α-tACS Lasts Up to 70 min after Stimulation. Front. Hum. Neurosci. 10, 245–245. doi:https://doi.org/10.3389/fnhum.2016.00245

Kasten, F. H., Duecker, K., Maack, M. C., Meiser, A., Herrmann, C. S., 2019. Integrating electric field modeling and neuroimaging to explain inter-individual variability of tACS effects. Nature Communications 10(1), 5427. doi:https://doi.org/10.1038/s41467-019-13417-6

Kasten, F. H., Herrmann, C. S., 2017. Transcranial Alternating Current Stimulation (tACS) Enhances Mental Rotation Performance during and after Stimulation. Front. Hum. Neurosci. 11(2). doi:https://doi.org/10.3389/fnhum.2017.00002

Kleiner, M., Brainard, D., Pelli, D., 2007. What’s new in Psychtoolbox-3? Perception 36, 1–16. doi:https://doi.org/10.1177/03010066070360S101

Krause, M. R., Vieira, P. G., Csorba, B. A., Pilly, P. K., Pack, C. C., 2019. Transcranial alternating current stimulation entrains single-neuron activity in the primate brain. Proceedings of the National Academy of Sciences 116(12), 5747. doi:https://doi.org/10.1073/pnas.1815958116

Kubicki, S., Herrmann, W. M., Fichte, K., Freund, G., 1979. Reflections on the topics: EEG frequency bands and regulation of vigilance. Pharmakopsychiatrie, Neuro-Psychopharmakologie 12(2), 237–245. doi:https://doi.org/10.1055/s-0028-1094615

Kuhlman, W. N., 1978. Functional topography of the human mu rhythm. Electroencephalogr. Clin. Neurophysiol. 44(1), 83–93. doi:https://doi.org/10.1016/0013-4694(78)90107-4

Larson, E., Taulu, S., 2018. Reducing Sensor Noise in MEG and EEG Recordings Using Oversampled Temporal Projection. IEEE Trans. Biomed. Eng. 65(5), 1002–1013. doi:https://doi.org/10.1109/tbme.2017.2734641

Narici, L., Portin, K., Salmelin, R., Hari, R., 1998. Responsiveness of Human Cortical Activity to Rhythmical Stimulation: A Three-Modality, Whole-Cortex Neuromagnetic Investigation. NeuroImage 7(3), 209–223. doi:https://doi.org/10.1006/nimg.1998.0323

Neuling, T., Wagner, S., Wolters, C. H., Zaehle, T., Herrmann, C. S., 2012. Finite-Element Model Predicts Current Density Distribution for Clinical Applications of tDCS and tACS. Frontiers in Psychiatry 3, 83. doi:https://doi.org/10.3389/fpsyt.2012.00083

Reinhart, R. M. G., Nguyen, J. A., 2019. Working memory revived in older adults by synchronizing rhythmic brain circuits. Nat. Neurosci. 22(5), 820–827. doi:http://doi.org/10.1038/s41593-019-0371-x

Romei, V., Gross, J., Thut, G., 2012. Sounds Reset Rhythms of Visual Cortex and Corresponding Human Visual Perception. Curr. Biol. 22(9), 807–813. doi:https://doi.org/10.1016/j.cub.2012.03.025

Simon, M., Schmidt, E. A., Kincses, W. E., Fritzsche, M., Bruns, A., Aufmuth, C., … Schrauf, M., 2011. EEG alpha spindle measures as indicators of driver fatigue under real traffic conditions. Clin. Neurophysiol. 122(6), 1168–1178. doi:https://doi.org/10.1016/j.clinph.2010.10.044

Stecher, H. I., Herrmann, C. S., 2018. Absence of Alpha-tACS Aftereffects in Darkness Reveals Importance of Taking Derivations of Stimulation Frequency and Individual Alpha Variability Into Account. Front. Psychol. 9(984). doi:https://doi.org/10.3389/fpsyg.2018.00984

Stecher, H. I., Pollok, T. M., Strüber, D., Sobotka, F., Herrmann, C. S., 2017. Ten Minutes of α-tACS and Ambient Illumination Independently Modulate EEG α-Power. Front. Hum. Neurosci. 11, 257. doi:https://doi.org/10.3389/fnhum.2017.00257

Takahashi, T., Kitazawa, S., 2017. Modulation of Illusory Reversal in Tactile Temporal Order by the Phase of Posterior α Rhythm. The Journal of Neuroscience 37(21), 5298–5308. doi:https://doi.org/10.1523/jneurosci.2899-15.2017

Taulu, S., Simola, J., Kajola, M., 2005. Applications of the signal space separation method. IEEE Transactions on Signal Processing 53(9), 3359–3372. doi:https://doi.org/10.1109/TSP.2005.853302

Vanni, S., Portin, K., Virsu, V., Hari, R., 1999. Mu rhythm modulation during changes of visual percepts. Neuroscience 91(1), 21–31. doi:https://doi.org/10.1016/S0306-4522(98)00521-1

Veniero, D., Benwell, C. S. Y., Ahrens, M. M., Thut, G., 2017. Inconsistent Effects of Parietal α-tACS on Pseudoneglect across Two Experiments: A Failed Internal Replication. Front. Psychol. 8(952). doi:https://doi.org/10.3389/fpsyg.2017.00952

Vossen, A., Gross, J., Thut, G., 2015. Alpha Power Increase After Transcranial Alternating Current Stimulation at Alpha Frequency (α-tACS) Reflects Plastic Changes Rather Than Entrainment. Brain Stimul. 8(3), 499–508. doi:https://doi.org/10.1016/j.brs.2014.12.004

Vosskuhl, J., Huster, R. J., Herrmann, C. S., 2015. Increase in short-term memory capacity induced by down-regulating individual theta frequency via transcranial alternating current stimulation. Front. Hum. Neurosci. 9, 257. doi:https://doi.org/10.3389/fnhum.2015.00257

Yokosawa, K., Murakami, Y., Sato, H., 2020. Appearance and modulation of a reactive temporal-lobe 8–10-Hz tau-rhythm. Neurosci. Res. 150, 44–50. doi:https://doi.org/10.1016/j.neures.2019.02.002

Zaehle, T., Rach, S., Herrmann, C. S., 2010. Transcranial Alternating Current Stimulation Enhances Individual Alpha Activity in Human EEG. PLoS One 5(11), e13766. doi:https://doi.org/10.1371/journal.pone.0013766

