## Supplementary material for "Transcranial alternating current stimulation affects several alpha components depending on their frequencies relative to the stimulation frequency": Supplementary_materials.pdf

### Supplementary Information

#### 1. Parameter settings for the fitting-oscillations-and-one-over-f (FOOOF) toolbox

The parameters of the FOOOF toolbox (Donoghue et al., 2020) were set as: peak width limits: [0.2, 1.5] or [0.2, 2.0] depending on the data; max number of peaks: 4; minimum peak height: 0 (default); peak threshold: 2.0 (default); and aperiodic mode: ‘fixed’. For the peak width limits, the values with a higher fitting accuracy were used. The functions of the PSDs were parameterised across a frequency range of 1.5 to 20.0 Hz.

#### 2. Frequency analysis with a low-frequency resolution

To confirm the validity of separating multiple alpha components with high-frequency resolution, we also performed the same frequency analysis as Kasten et al. (2019) with low-frequency resolution. After removing the artefacts, continuous signals were cut into 2 s segments. Segments containing the timing of the rotation of the visual stimuli and button responses were excluded. Power spectral densities (PSDs) were calculated using a multitaper fast Fourier transform (FFT; 2 s zero-padding, frequency resolution = 0.5 Hz, interval of frequency bins = 0.25 Hz) on the first 260 artefact-free segments in each period. Note that zero-padding does not improve the frequency resolution of the FFT, but increases only the point-to-point interval in the resulting PSDs because the length of an actual signal does not change (Ciaccio et al., 2012).

The PSDs of the sensor signals before and after stimulation were calculated for each stimulation condition, and the averages of all gradiometers and participants are shown in Supplementary Figure 2. All PSDs were aligned with the participants’ individual alpha frequency (IAF) estimated from the pre-stimulation period. There were significant power changes from pre- to post-stimulation in the frequency band lower than the IAF and 5–10 Hz higher than the IAF in both conditions in the sensor analysis (Wilcoxon signed-rank test, $p < 0.05$, Bonferroni-corrected). Peak alpha frequency decreased in both sham-stimulation and transcranial alternating current stimulation (tACS) conditions ($t$ (34) = 4.96, $p < 0.01$, Cohen's $d = 0.64$; $t$ (34) = 2.69, $p = 0.02$, Cohen's $d = 0.54$, paired $t$-test). These findings are similar to those of a previous study that reported a power increase and frequency decrease in the alpha peak during a task (Benwell et al., 2019). Importantly, Benwell et al. (2019) suggested that these changes are caused by different alpha components.

In contrast, in the source domain, we found no significant clusters among participants in either condition, which is inconsistent with Kasten et al. (2019). They reported that the increase in alpha power in the sham-stimulation group was confined to a few occipital, posterior-parietal, and temporal regions, whereas the increase in the tACS group spread over a wide range of cortical areas, including occipital-parietal, temporal, and frontal regions. As a result, the aftereffects of tACS, defined by the interaction between period (pre- vs. post-stimulation) and stimulation condition (tACS vs. sham), extended over a wide area of the cortex, including the frontal region not covered by electrode montage (Kasten et al., 2019). In contrast, in the current study, while a significant power increase around the IAF from pre- to post-stimulation in sensor analysis was observed, there was no significant increase in the source space from pre- to post-stimulation in both conditions. This is probably because the source of the alpha power change between pre- and post-stimulation varied from participant to participant. Other studies have suggested that multiple components exist in the alpha frequency band (Barzegaran et al., 2017; Chiang et al., 2011; Takahashi and Kitazawa, 2017). Based on these observations, we attempted to decompose the multiple alpha components and evaluate them individually.

### 3. Clustering analysis based on spatial topographic patterns of two alpha peaks

A typical structure of resting-state alpha rhythms originates from the occipital-parietal and/or occipital-temporal regions (Barzegaran et al., 2017). Therefore, we performed hierarchical cluster analysis using the Ward method with the linkage function of SciPy, a Python library, to classify all participants into several groups based on the characteristics of the two topographic maps of an individual's alpha peaks. Silhouette analysis was also performed to ensure that all silhouette values were not negative, and that the maximum value for each cluster member exceeded the average silhouette value for all participants. The silhouette value is a measure of the degree of cohesion within a cluster and the degree of divergence from other clusters, ranging from 1 at the maximum separation performance to −1 at the worst (Rousseeuw, 1987).

All the silhouette values were positive when the number of clusters was set to two. We termed these two clusters the temporal/occipital group and temporo-occipital/parietal group based on the spatial characteristics of the alpha components. The mean silhouette value was 0.17, with a maximum value of 0.19 in the temporal/occipital group and 0.38 in the temporo-occipital/parietal group. When the number of clusters was set to three or four, the average silhouette coefficients were -0.08 or -0.05, respectively.

Although this classification was based on topographic patterns of alpha peaks, it could have been based on other criteria. Since the peak frequencies of the temporal, occipital, and parietal components are in ascending order, and the occipital component is evident for all participants (See Supplementary Table 1), we also successfully classified participants by considering whether the peak frequency reflecting the occipital component was higher or lower than the other peak. Because this method does not change the classification of the participants, our classification of participants would be robust.

#### 4. Subjective evaluation of drowsiness

Immediately after the magnetoencephalography measurement, the participants gave subjective evaluations of their drowsiness during the experiment. Specifically, the following five sentences were presented in Japanese, and participants were asked to mark the appropriate one: (i) “I was not sleepy at all from the beginning to the end”; (ii) “I became sleepy towards the second half of the session”; (iii) “I became less sleepy over the second half of the session”; (iv) “I was sleepy the whole time”; or (v) “I fell asleep during the session”. The results are presented in Supplementary Figure 3. Sixty-nine percent of the participants felt drowsy during the experiment, and more than half of them became drowsy over time.

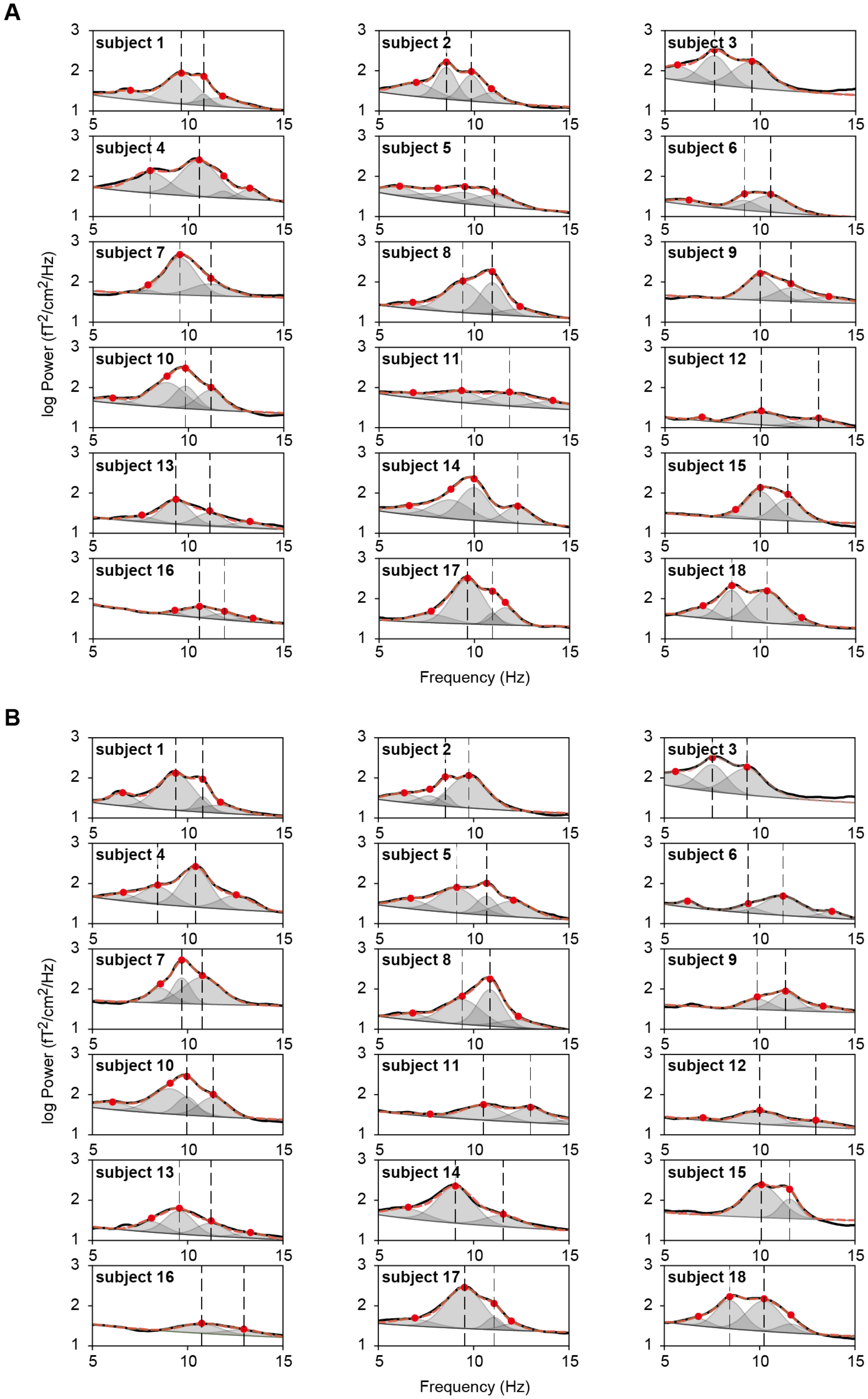
A
subject 1
subject 2
subject 3
subject 4
subject 5
subject 6
subject 7
subject 8
subject 9
subject 10
subject 11
subject 12
subject 13
subject 14
subject 15
subject 16
subject 17
subject 18
log Power (fT2/cm2/Hz)
Frequency (Hz)
B
subject 1
subject 2
subject 3
subject 4
subject 5
subject 6
subject 7
subject 8
subject 9
subject 10
subject 11
subject 12
subject 13
subject 14
subject 15
subject 16
subject 17
subject 18
log Power (fT2/cm2/Hz)
Frequency (Hz)

Supplementary Figure 1. Individual FOOOF fitting results. Each panel represents the PSD of an individual participant in the sham-stimulation condition (A) and in the tACS condition (B). The black line indicates an original PSD, and the dashed red line indicates the sum of multiple Gaussian fittings. Red points represent a peak of each Gaussian distribution. The frequencies of the two largest products of a height of the periodic component and the height of the Gaussian distribution were defined as the peak alpha frequencies for that participant.

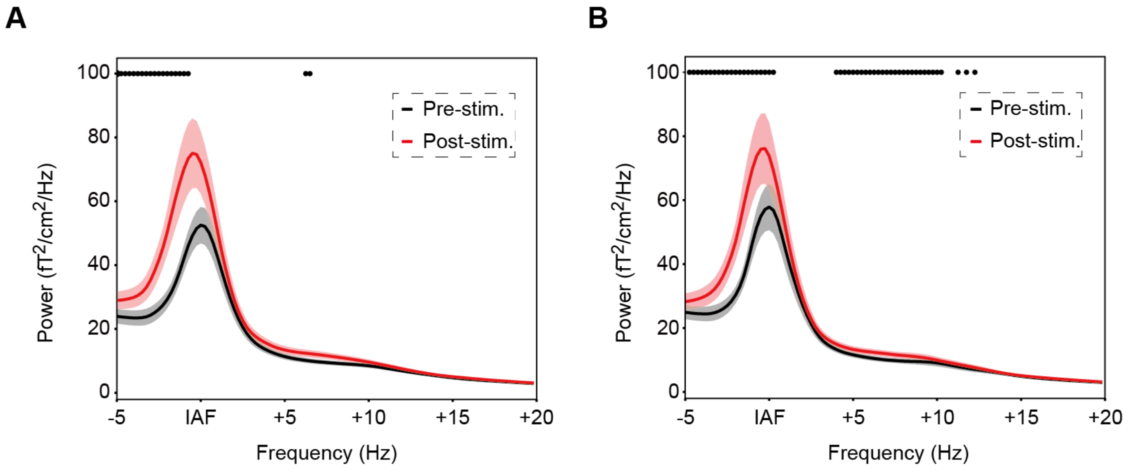


Supplementary Figure 2. Averaged PSDs of all participants (A) in the sham-stimulation condition and (B) in the tACS condition analysed with low-frequency resolution (0.5 Hz). Black dots show significant differences between pre- and post-stimulation (Wilcoxon signed-rank test, $p < 0.05$, Bonferroni-corrected). The shaded areas represent the standard errors. In both the sham-stimulation and tACS conditions, the power increase and the peak frequency decrease were observed. No significant difference was found between stimulation conditions.

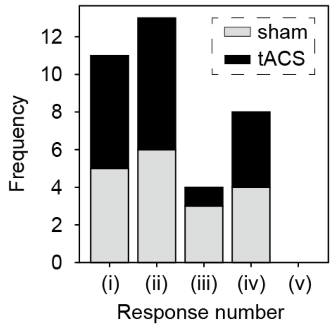

Supplementary Figure 3. The results of the subjective evaluation of drowsiness collected after each tACS experiment. The following five sentences were presented in Japanese: (i) “I was not sleepy at all from the beginning to the end”; (ii) “I became sleepy towards the second half of the session”; (iii) “I became less sleepy over the second half of the session”; (iv) “I was sleepy the whole time”; or (v) “I fell asleep during the session”. We found 69% of participants felt drowsy during the experiment, and moreover, many of them became drowsier over time (ii).

Supplementary Table 1. Dominant alpha frequencies of two groups of participants.

| Temporal/Occipital group | | | Temporo-Occipital/Parietal group | | |
|---|---|---|---|---|---|
| **subject ID** | **Low peak** Temporal comp. | **High peak** Occipital comp. | **subject ID** | **Low peak** Temporo-Occipital comp. | **High peak** Parietal comp. |
| 2 | 8.5 Hz | 9.7 Hz | 1 | 9.4 Hz | 10.8 Hz |
| 3 | 7.5 Hz | 9.3 Hz | 9 | 9.9 Hz | 11.4 Hz |
| 4 | 8.4 Hz | 10.4 Hz | 10 | 10.0 Hz | 11.3 Hz |
| 5 | 9.1 Hz | 10.7 Hz | 11 | 10.5 Hz | 13.0 Hz |
| 6 | 9.4 Hz | 11.2 Hz | 12 | 10.0 Hz | 12.9 Hz |
| 7 | 9.7 Hz | 10.8 Hz | 13 | 9.6 Hz | 11.2 Hz |
| 8 | 9.4 Hz | 10.9 Hz | 14 | 9.0 Hz | 11.6 Hz |
| 18 | 8.4 Hz | 10.2 Hz | 15 | 10.1 Hz | 11.6 Hz |
| | | | 16 | 10.7 Hz | 12.9 Hz |
| | | | 17 | 9.5 Hz | 11.1 Hz |

### References


Barzegaran, E., Vildavski, V. Y., Knyazeva, M. G., 2017. Fine Structure of Posterior Alpha Rhythm in Human EEG: Frequency Components, Their Cortical Sources, and Temporal Behavior. Sci. Rep. 7(1), 8249. doi:https://doi.org/10.1038/s41598-017-08421-z

Benwell, C. S. Y., London, R. E., Tagliabue, C. F., Veniero, D., Gross, J., Keitel, C., Thut, G., 2019. Frequency and power of human alpha oscillations drift systematically with time-on-task. NeuroImage 192, 101-114. doi:https://doi.org/10.1016/j.neuroimage.2019.02.067

Chiang, A. K., Rennie, C. J., Robinson, P. A., van Albada, S. J., Kerr, C. C., 2011. Age trends and sex differences of alpha rhythms including split alpha peaks. Clin. Neurophysiol. 122(8), 1505-1517. doi:https://doi.org/10.1016/j.clinph.2011.01.040

Ciaccio, E. J., Biviano, A. B., Whang, W., Garan, H., 2012. Improved frequency resolution for characterization of complex fractionated atrial electrograms. Biomed. Eng. Online 11(1), 17. doi:https://doi.org/10.1186/1475-925X-11-17

Donoghue, T., Haller, M., Peterson, E. J., Varma, P., Sebastian, P., Gao, R., . . . Voytek, B., 2020. Parameterizing neural power spectra into periodic and aperiodic components. Nat. Neurosci. 23(12), 1655-1665. doi:https://doi.org/10.1038/s41593-020-00744-x

Kasten, F. H., Duecker, K., Maack, M. C., Meiser, A., Herrmann, C. S., 2019. Integrating electric field modeling and neuroimaging to explain inter-individual variability of tACS effects. Nature Communications 10(1), 5427. doi:https://doi.org/10.1038/s41467-019-13417-6

Rousseeuw, P. J., 1987. Silhouettes: A graphical aid to the interpretation and validation of cluster analysis. Journal of Computational and Applied Mathematics 20, 53-65. doi:https://doi.org/10.1016/0377-0427(87)90125-7

Takahashi, T., Kitazawa, S., 2017. Modulation of Illusory Reversal in Tactile Temporal Order by the Phase of Posterior α Rhythm. The Journal of Neuroscience 37(21), 5298-5308. doi:https://doi.org/10.1523/jneurosci.2899-15.2017
